# A survey of Rare Disease awareness among healthcare professionals and researchers in India

**DOI:** 10.1101/2023.03.31.534989

**Authors:** Laxmikant Vashishta, Purna Bapat, Yashodhara Bhattacharya, Mohua Chakraborty Choudhury, Narendra Chirmule, Susan D’Costa, Shilpa Jindani, Manohar Rao, Smritie Sheth

## Abstract

Rare diseases (RDs) are diseases that occur infrequently and affect a small fraction of the population. Although these diseases individually affect small number of people, together they affect 400 million people globally at any given time. In India, where resources are scarce, healthcare infrastructure and policy framework are focused on mitigating diseases that affect many people. Further, the level of RD awareness among healthcare professionals, researchers, and general public is considerably low. As a result, many cases of RDs remain unreported, undiagnosed, and untreated. To frame policies regarding RDs, it is crucial to understand the current level of RD awareness among healthcare professional and researchers, as they are key stakeholders in diagnosis, treatment, policy making, and drug development. We conducted an exploratory survey to understand the current level of RD awareness among healthcare professionals and researchers based on identification of an RD, time for diagnosis, treatment options, and relationship with family history and geographic location. We noted that our respondents have considerably low level of RD awareness. They correctly identified the importance of family history but failed to realize the association with geographic location. After presenting the survey findings, we have made recommendation to improve RD awareness in India. Our findings will be helpful to design awareness campaigns and frame relevant policies.

## Introduction

Rare diseases (RDs) is an umbrella term used to define diseases that occur infrequently and affect a relatively small fraction of the population [1]. These conditions are often serious, life-debilitating, affect multiple organs, and have a severe impact on the lives of patients and their families. RDs are a heterogeneous group of diseases, and around 7000-8000 different RDs have been identified globally (ref). Although individually each RD affects a few people, together they affect 400 million people globally at any point in time, accounting for 4% of the total world population [2]. However, there is no universal definition of an RD. A study identified 296 definitions from 1109 organizations [3]. Different countries have adopted different prevalence thresholds for classifying RDs. For example, in the USA, RDs are defined diseases or conditions that affect fewer than 200,000 people at any point in time, whereas in the EU, RDs are diseases that affect no more than 1 in 2000 people.

As any RD individually affects a very small number of people who are geographically dispersed, most healthcare professionals (HCPs) are not aware of such diseases. There is a lack of knowledge and understanding of RDs, which is because there is no significant research about RDs. In addition, the pharmaceutical industry does not find it profitable to develop drugs for such a small patient size. Thus, RDs do not typically feature in the healthcare policy of most countries. Despite these challenges, as a result of huge patient advocacy movements intended to draw attention towards RDs, the Rare Drug Act (RDA) [4] was passed in the United States 1983, which provided various incentives to pharmaceutical companies to develop drugs for RDs. RDA has proven to be a game changer in the US; it has driven the development of more than 600 approved drugs for RDs. This number was less than 10 across different therapeutic areas and disease categories in the decade before the RDA was passed. This also led to an increase in awareness and interest among the public, healthcare professionals, and researchers. The success of RDA in the US inspired patient advocacy groups in other countries to drive awareness campaigns and advocate for legislative changes to address the unmet needs of RD patients, especially in the high-income countries such as the EU, Japan, and China.

Healthcare policies in India, where resources are scarce, are focused on mitigating diseases that affect large populations. Policies for RDs have not figured in any government agenda and policies until recently. The revised National Policy for Treatment of Rare Diseases (NPTRD) [5] is the first policy in India dedicated to addressing concerns of the RD community. It was first released in 2017, but it was subsequently put in abeyance because of major implementation challenges [6, 7]. The New Drugs and Clinical Trials Rule [8], conceptualized in 2019, also had some incentives for the approval and marketing of Rare drugs. These policies acknowledge the lack of awareness, epidemiological data, and knowledge resources on RDs in India. Only 450 RDs have currently been identified in India, and most RDs go undiagnosed and unrecorded. Patients who have received a diagnosis have reported facing many difficulties and many years of waiting before receiving a correct diagnosis [9]. They also reported a huge lack of awareness among HCPs. To strengthen the RD ecosystem of the country, it is imperative that there is increased awareness among HCPs and researchers. Lack of awareness was cited as a major roadblock in indigenous development of Rare medical products [10]. Increased awareness among HCPs will help to address some of the major challenges faced by RD patients, such as delayed diagnosis, misdiagnosis, and improper treatment. Moreover, to improve diagnosis and management, it is imperative to have more research focused on understanding the natural history, epidemiology, prevalence among the local population, elucidating etiological factors, and drug discovery/development of RDs. It is thus important to encourage researchers to develop an interest in RD-related research. HCPs and researchers also play a major role in generating awareness among other stakeholders through the dissemination of knowledge via various platforms. Furthermore, through their highly influential professional medical associations and research groups, both HCPs and researchers can play a big role in influencing government, policymakers, and the pharma industry to address the needs of the RD community.

Considering the important role of awareness in mitigating RDs, it is important to assess the level of RD awareness among HCPs and researchers. Similar studies in other countries have proved to be valuable in designing strategies to increase RD awareness among the concerned groups [11]. The study is rightly timed as it coincides with the release of the NPTRD and the outcome of the study will help in developing its implementation strategy with a focus on increasing awareness among the two stakeholder groups. To understand the awareness and interest about RDs among HCPs and biology researchers in India, we conducted an exploratory survey. The motivation of the survey was to collect preliminary data from HCPs and researchers on the level of their awareness about topics such as i) recent developments in diagnosis, treatment, and clinical trials, ii) geographies associated with RDs, and iii) societies and associations for supporting RDs. The goal is to extract sufficient insights to be able to develop a study to evaluate the current state of knowledge of RDs in India.

## Methods

### Survey

We conducted an exploratory survey to understand various aspects of rare diseases in India. The survey was not aimed to determine statistically significant results, rather obtain a preliminary understanding of the trends. The main objective of the survey was to understand the awareness of rare diseases in India among HCPs and Researchers. The responses were deemed sufficient as a representative group of HCPs and Researchers were surveyed to evaluate the awareness of rare diseases and their prevalence in India. These preliminary findings will be used to develop several hypotheses for a future study.

Two separate questionnaires were developed for this survey: one for HCPs and another for researchers. The questionnaire for HCPs had 22 questions, whereas the one for researchers had 22. The questionnaires are provided in Appendices 1 and 2. The questions were framed based on the following points: i) awareness of specific RDs, ii) diagnoses, iii) genetic testing and prevention, iv) treatment, v) molecular basis of the RD, vi) sources of information, and vii) demographics

The research team reached out to HCPs and researchers in India. The selected respondents included practicing doctors/physicians, nurses, genetic counselors, nurses and researchers in biomedical sciences. The survey was distributed by the research team through email, social media (LinkedIn™, Facebook™, Whatsapp™), and personal communication. There were a total of 117 respondents: 52 HCPs and 65 researchers. Given the process of data collection, we anticipated that there was minimum bias in results, and the respondents were representative of the groups from which the information was intended.

The responses to the survey were recorded using Google forms, and the data was downloaded into an Excel spreadsheet. Prior to analysis of the results, the data was curated: invalid data were excluded, and repetitive data were combined. For example, i) blank cells were deleted and same or similar words (e.g. gene, gene therapy, genomic) were combined; ii) repeated words were omitted; iii) spelling errors were corrected; iv) punctuation errors were fixed)

## Results

### Demographics of the survey respondents

A total of 117 respondents participated in the survey. The HCPs (24) were mainly physicians (MBBS/MD), genetic counsellors, nurses, and other personnel involved in point of care diagnosis and treatment of RDs. The researchers (76) were either Master’s or PhD holders in biological sciences, as they were approached via the biological sciences institutes to which they were affiliated. The demographic spread of survey respondents is presented in Table 1.

**Table 1.** Demographics of the survey respondents. The columns represent educational qualifications, whereas the rows represent current professions.

| **Self – description** | **MBBS/MD** | **Genetic**<br>**Counselling** | **Master’s** | **PhD** | **Grand**<br>**Total** |
| --- | --- | --- | --- | --- | --- |
| **Doctor(physician)** | 23 | 1 |  | 11 | 35 |
| **Genetic Counselor** | 1 | 9 |  | 1 | 11 |
| **Genetic**<br>**Technologist** |  |  | 1 |  | 1 |
| **Researchers** |  |  | 41 | 24 | 65 |
| **Others** |  | 1 | 3 | 1 | 5 |
| **Grand Total** | 24 | 11 | 45 | 37 | 117 |

#### 2.1 Awareness of RDs among HCPs and researchers

The survey participants were asked to mention one or more RDs that they were most aware of. They might have either worked on it as a researcher or diagnosed and/or treated it as an HCP. The data collected for this question was represented and analyzed as a word cloud. In this word cloud, the RDs were written in font sizes proportional to the number of times they were mentioned by the respondents. The data from HCPs and researchers were represented in different word clouds to understand the differences in their responses (**Figures 1a and 1b**). Figures 1a and 1b show that the RDs identified by HCPs and researchers were different. The figure shows that HCPs identified Huntington’s disease, Spinal Muscular Atrophy, Alkaptonuria, Thalassemia, Klinefelter Syndrome, Duchenne Muscular Dystrophy, Marfan Syndrome and Neurofibromatosis among other Rare diseases while Researchers identified Hemophilia, Alzheimer’s Disease, Spinal Muscular Atrophy, Huntington’s Disease, Amyotrophic Lateral Sclerosis (ALS) and Niemann Pick Syndrome among other Rare diseases. These listed Rare diseases were identified more times than other Rare diseases by both HCPs and Researchers. Some of the diseases, e.g., Huntington’s disease and Spinal Muscular Atrophy (SMA), were named by both HCPs and researchers.

**Figure 1a.**
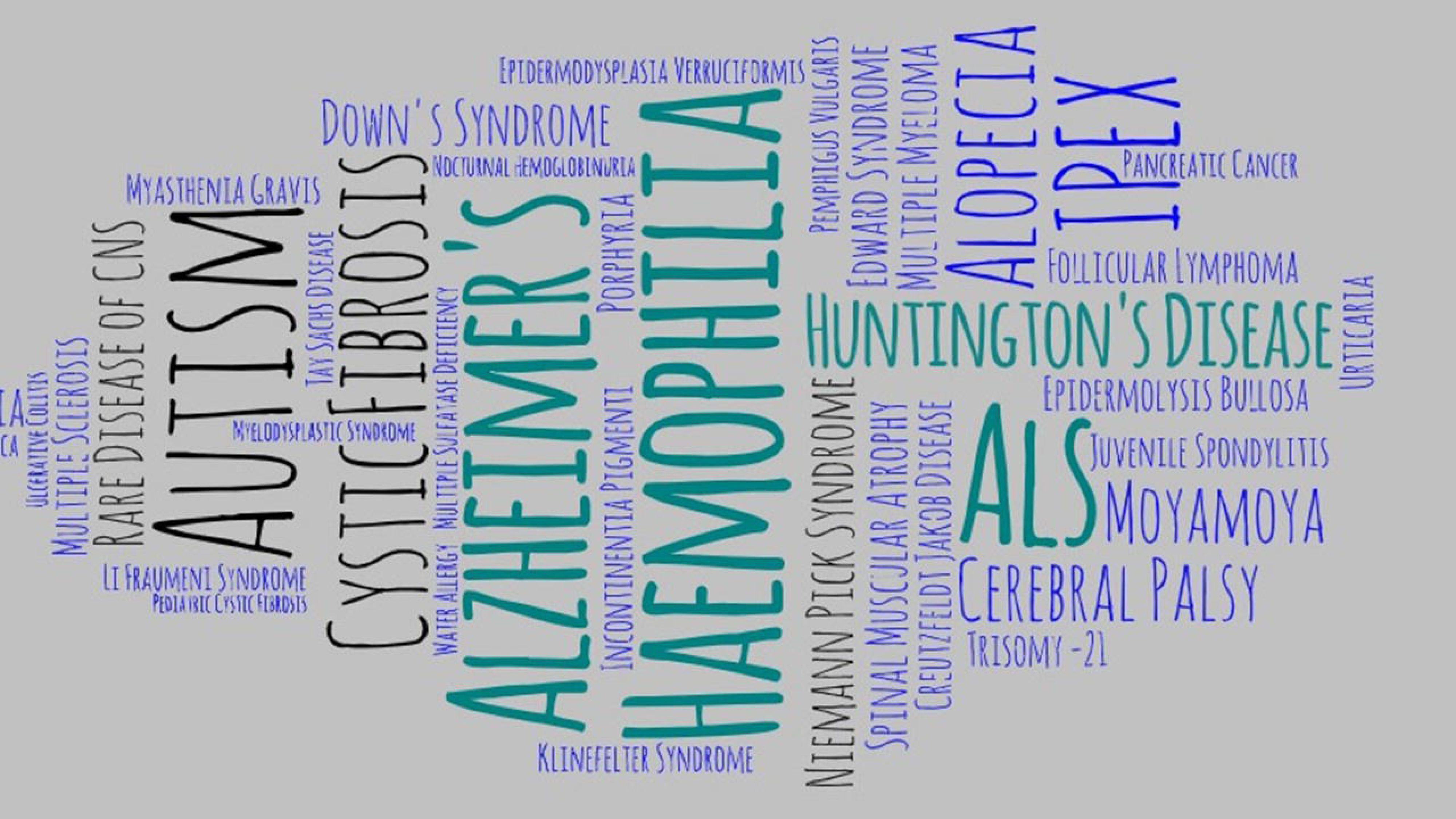
Word Cloud: HCPs. This word cloud represents different rare diseases selected or named by HCPs. The size and color of the fonts represent the number of times that disease was selected or named.

**Figure 1b.**
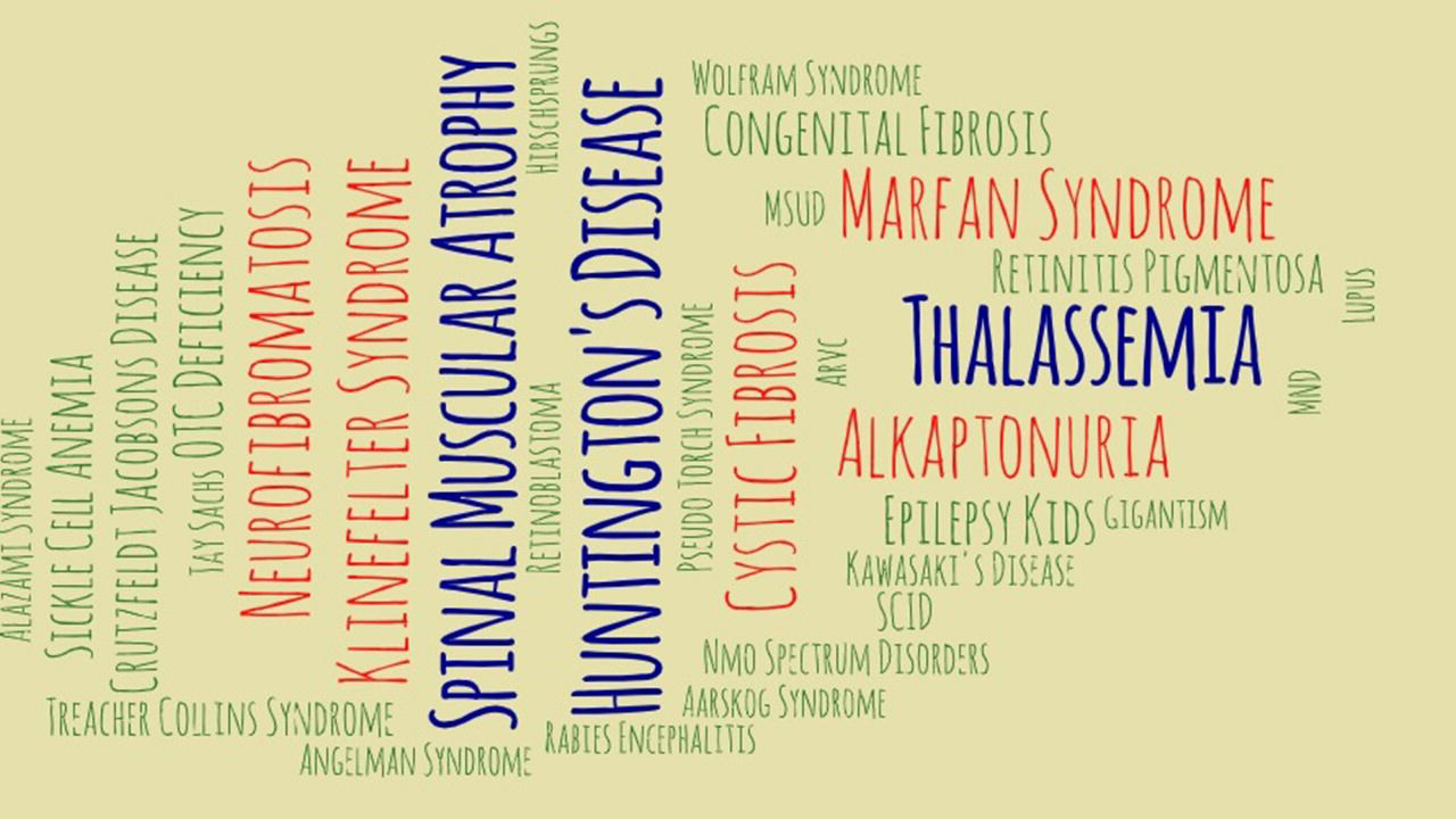
Word Cloud: Researchers. This word cloud represents different rare diseases selected or named by researchers. The size and color of the fonts represent the number of times that disease was selected or named.

#### 2.2 Time to diagnosis of RDs

Survey respondents were asked on average how long it takes for the RD they have mentioned to be diagnosed after the patient first presents symptoms. Seventeen (34.6%) HCPs and 16 (27.5%) researchers believed that it takes less than a year to diagnose the RD they had mentioned in the previous question. HCPs identified the following diseases as the ones that take more than a year to diagnose: Aarskog Syndrome, Alazani Syndrome, Alkaptonuria, Cystic Fibrosis, Duchenne Muscular Dystrophy, Gigantism, Hirschsprung disease, Klinefelter Syndrome, Maple Syrup Urine Disease, NMO Spectrum Disorders, OTC Deficiency, Pseudo Torch Syndrome, Rabies Encephalitis, Retinoblastoma, Severe Combined Immune Deficiency (SCID), Spinal Muscular Atrophy (SMA), and Treacher Collins Syndrome. Researchers, on the contrary, identified the following diseases as ones that require less than a year to diagnose: Down’s Syndrome, Edward’s Syndrome, Epidermolysis Bullosa, Immunodysregulation polyendocrinopathy enteropathy X-linked (IPEX) Syndrome, Li-Fraumeni Syndrome, Multiple Sulfatase Deficiency, Pemphigus Vulgaris, Persistent Cloaca, Progeria, Hemophilia, SMA, Tay Sachs Disease, and Ulcerative Colitis. Twenty-Three percent HCPs and 18% researchers mentioned that it takes 1-3 years to diagnose an RD of their choice. In this category, HCPs identified Angelman Syndrome, Cystic Fibrosis, Huntington’s Disease, Marfan Syndrome, Neurofibromatosis, Retinitis Pigmentosa, Sickle Cell Anemia, and Thalassemia. Researchers identified Amyotrophic Lateral Sclerosis, Autism, Follicular Lymphoma, Huntington’s Disease, Incontinentia Pigmenti, Myasthenia Gravis, Pancreatic Cancer, and Pediatric Cystic Fibrosis. Overall, 15% HCPs and 23% researchers mentioned that it takes more than 5 years to diagnose the disease. HCPs identified DMD, Huntington’s Disease, Lupus, Marfan Syndrome, Neurofibromatosis, and Thalassemia in this category (**Figure 2**).

**Figure 2.**
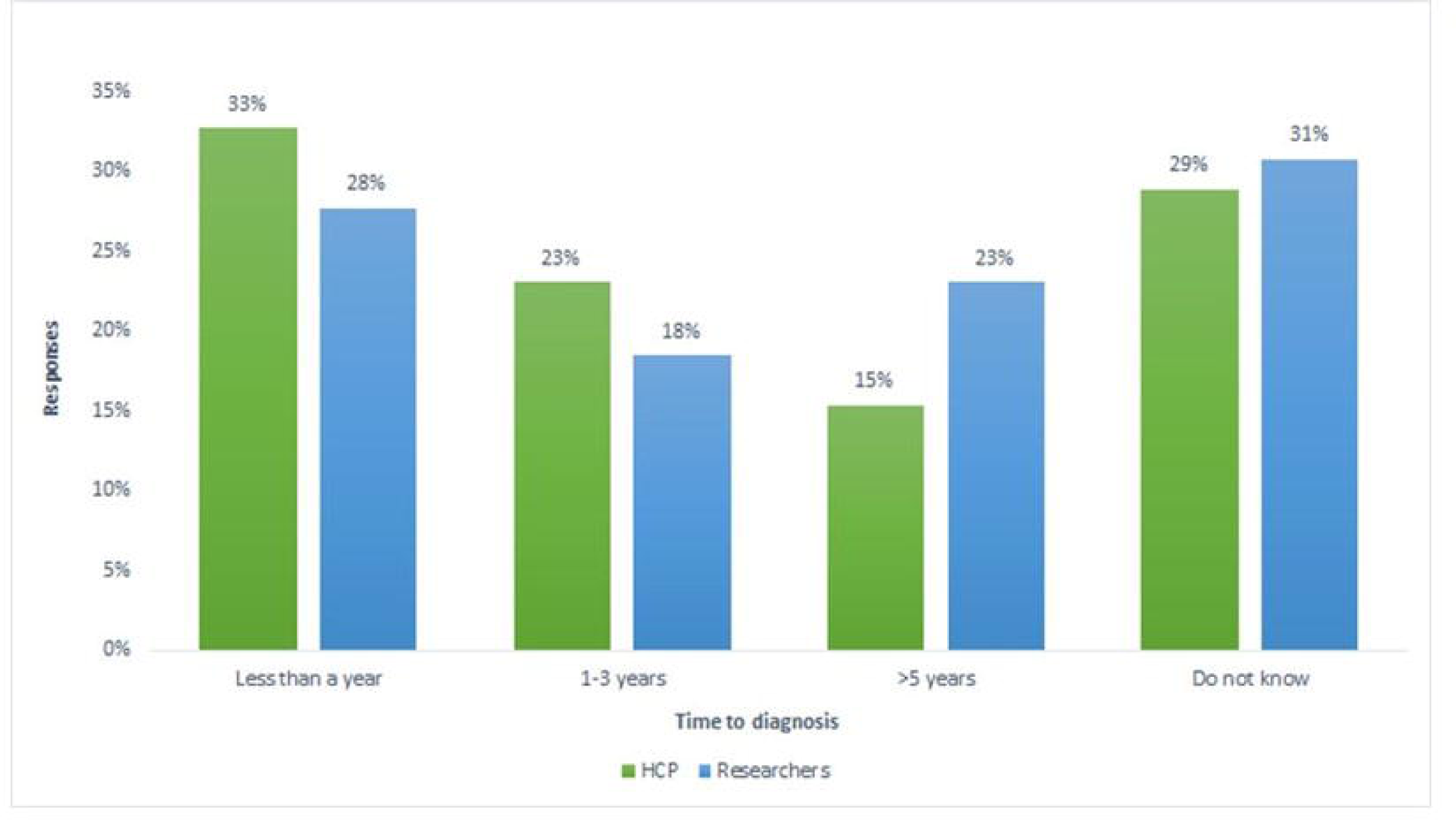
Time required to diagnose an RD. The bar graph represents opinions of survey respondents on the time required to diagnose the RD they have mentioned.

#### 2.3 Diagnosis and treatment options for patients with RDs

HCPs were asked to select the treatment options that were most effective in treating the RD that they chose in in a previous question. The responses collected were analyzed using a mosaic plot (**Figure 3**). This plot shows the comparison of the opinions given by various HCPs about different options for treating RDs. In the survey, many respondents chose more than one options; therefore, to account for such responses, a separate category of “multiple selections” was created. Similarly, some respondents provided answers that were not in the list; these were placed in the “others” category. This analysis revealed that out of 52 HCP respondents, physician-HCP’s [8 (33%)]were of the opinion that symptomatic management is the most effective way of treating RDs. Another 5 physician-HCP’s (20.8%) believed that gene/cell therapy is the most effective way and 4 (16.6%) said that other treatment options could be beneficial. On the contrary, the non-physician-HCPs chose biologics (6 of 13; 46%) as the treatment option and remaining ones chose gene/cell therapy (2 of 13; 15%). Out of the 11 genetic counselors, 3 (27%) chose gene therapy as the treatment option. Further, individual treatment, symptom management, multiple selection, and other categories received equal responses.

**Figure 3.**
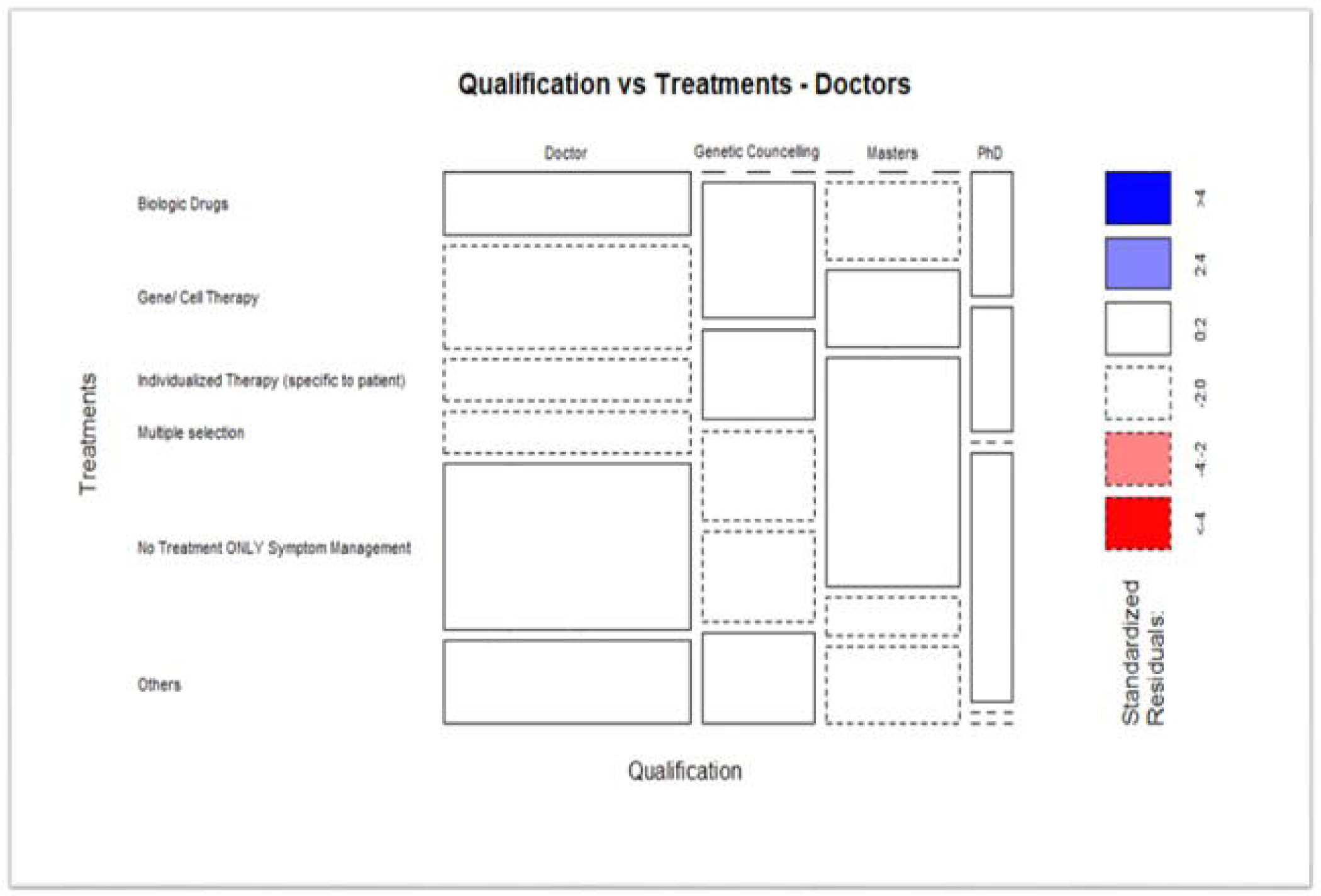
Treatment options for RDs versus qualifications of respondents.

#### 2.4 Sources of RD-related information

HCPs and researchers were asked how they kept themselves updated about the recent developments in RD research. For this question, 26 (50%) respondents selected more than one sources for receiving information about RDs. To account for responses that involved more than one options, a separate category of “multiple selections” was created for data visualization. The analyzed data are presented in Figure 4a. Out of 52 HCPs, 14 (26.92%) chose research papers across all affiliations, signifying that research papers were the most preferred source of keeping oneself updated. In addition to research papers, HCPs chose other options as means to keep themselves updated on RDs. Within the multiple selections category, 25 out of 26 respondents selected research papers. This was followed by social media 6 (11.53%) and print media 4 (7.69%). In addition to the absolute numbers, we checked whether the affiliation of the respondents is associated with their means of obtaining information on RDs. We noted that HCPs from teaching hospitals prefer research papers (3 out of 7, 42.85%) more than the HCPs from other affiliations. Subsequently, HCPs from government hospitals (3 out of 8, 37.5%), industry (4 out of 16, 25%), private hospitals (2 out of 10, 20%), and private practice (2 out of 10, 20%) prefer research papers over other means.

**Figure 4.**
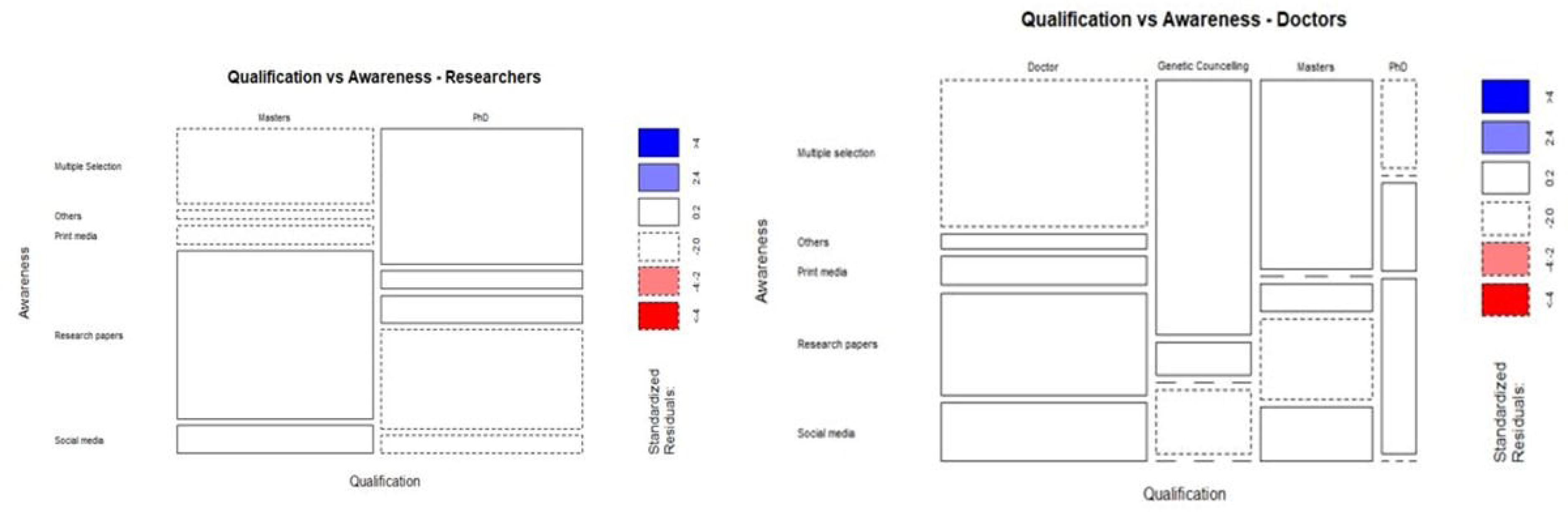
Association between qualifications of respondents and their awareness of RDs. (a) HCPs (b) researchers. Mosaic plots: Mosaic plot is a method of graphical visualization of data. It provides a visual comparison between different attributes. A mosaic plot represents the cell frequencies of the cross-tabulation. The width of the columns in the plot is proportional to the number of corresponding observations. A Mosaic Plot for 2 variables gives a visual representation of the relationship between 2 categorical variables. The width of the columns is proportional to the number of observations for each level within the variable on the X-axis. The vertical length of the bars is proportional to the number of observations in the second variable within each level of the first variable. The color of the plot shows the amount of deviation of the observed frequency from the expected frequency. The style of the border tells whether the deviation is positive or negative. If the residuals are highly significant, we can say that there is a high association between the levels.

**Figure 5.** Treatments available for RDs.

Similarly, in the researcher’s survey (**Figure 4b**), 29 out of 65 respondents (44%) chose research papers as a source of keeping themselves acquainted with updates on RDs. This was followed by multiple selections, which accounted for 23 out of 65 (35%), and print media and social media, accounted for 5 out of 65 (7%) each.

#### 2.5 Geographic distribution

HCPs and researchers were asked if the RD of their choice was endemic to any region of India. Most participants did not believe or did not know if RDs have an association with a geographic region. Among the HCPs, 50% did not believe and 30.8% did not know if RDs have an association with a geographic region. Among the researchers, 37% did not believe and 40% did not know if RDs have an association with a geographic region. As the answers to this question did not indicate any association of RDs to a geographic region, we inferred that both HCPs and researchers are unaware of this aspect of RDs.

#### 2.6 Importance of family history

All respondents were asked if family history is significant in identifying the RD of their choice. Most of the respondents (71%) across both categories responded positively, indicating that both HCPs and researchers acknowledge the role of genetics in RDs.

## Discussion

In this study, we surveyed HCPs and researchers, as they are the important players in the RD ecosystem. They are involved in not just providing treatment but also in research, diagnosis, drug development, and policy making. Considering their important role, we assessed their level of awareness regarding diagnosis, treatment, and counseling of RDs. The survey was shared with ∼1700 HCPs and researchers through direct emails, institutional outreach, social media (Linkedin™, Facebook™, and WhatsApp™), and professional network outreach. Out of the people who have seen this survey, only ∼1% people responded to the survey. This response rate shows a limited interest and/or awareness about RDs in India. The respondents of this survey had diverse backgrounds in terms of specialization area, qualification, and geographic location. Thus, the set of respondents was fairly diverse, which helped to reduce the bias.

### 3.1 Awareness of RDs

RDs is a collective term, and it represents a heterogeneous group of diseases. In our survey, HCP and Researchers identified a different RDs belonging to different disease categories, which rightly reflects the heterogenous nature of RDs. For example, among the blood-borne diseases, HCPs identified Thalassemia and Sickle cell anemia, and researchers identified Hemophilia. Among the diseases related to the brain and CNS, HCPs identified Spinal Muscular Atrophy (SMA), Huntington’s disease, Neuromyelitis Optica, and Researchers identified Alzheimer’s disease, Amyotrophic Lateral Sclerosis, Multiple Sclerosis in addition to SMA and Huntington’s disease. Among the metabolic disorders, HCPs identified Alkaptonuria, and researchers identified Niemann pick disease. The diseases identified by the respondents were cross-checked with available data and resources, which revealed that these diseases are indeed Rare in India. The definition of RD in policy documents in India continue be clarified and there is limited information on prevalence data for most diseases. Most diseases identified in our study by the HCP and researchers are rare in India, as indicated by the hospital-based and epidemiology studies for individual RDs (Table 2). However, many[10] of these diseases are among the more common RDs. For example, many HCPs and researchers have identified some of the blood disorders as RDs, which are suspected to be not rare in some parts of India due to their high prevalence among some indigenous communities. [12, 13]

**Table 2.** Summary of top RDs identified by HCPs and researchers and their identification as RDs in India in other sources.

| Top Diseases identified by HCP and Researchers | HCP | Researchers | Identified as Rare in India in other sources |
| --- | --- | --- | --- |
| Hemophilia | Yes | No | No (ref 3,4) |
| SMA | Yes | No | Yes (ref 1, 2) |
| DMD | Yes | No | No |
| CF | Yes | Yes | Yes (ref 1, 2) |
| ALS | Yes | Yes | Yes (ref 1,2) |
| Alzheimer's | Yes | Yes | Yes (ref 1,2) |

### 3.2 Time for diagnosis

RDs are highly diverse in nature, and this is reflected in the differences in opinions of survey respondents regarding the time for diagnosis of RDs. It has been widely reported that it takes 7 years on an average to diagnose a RD in developed countries like the USA [10]. In India as well, the diagnostic odyssey has been widely reported for RDs. In addition, many cases remain undiagnosed, misdiagnosed, or take many years and multiple visits to several doctors before arriving at a correct diagnosis. In this survey, we see a lot of variation in responses regarding time to diagnosis: from less than 1 year to more than 5 years. Moreover, 29% of HCPs and 31% of researchers mentioned that they do not know anything about the time required for the diagnose a RD. This shows a poor level of awareness among these two communities in India. Further, 56% HCPs and 46% researchers mentioned that the RD they chose in a previous question requires less than 3 years for diagnosis.

This is contradictory to expert beliefs and findings from other studies [10] which indicate that RDs take more than 7 years for diagnosis in India, and in many cases, they remain undiagnosed. The NPRD appropriately raises concerns regarding the delay in diagnosis of RDs. Thus, the response that was received in our survey from a considerable number of respondents may be attributed to their lack of awareness about RDs. However, this response should not be generalized to all RDs, as the respondents were asked to select an RD in a previous question and the response regarding time to diagnosis was directed to that particular RD. The word cloud shows that Hemophilia, Autism, ALS, Alzheimer’s, Huntington’s Disease were among the most chosen RDs by the respondents. Some of the diseases that were selected by most of the participants do have a shorter diagnosis timeline. For example, Hemophilia, Autism, and Thalassemia have more evolved diagnosis protocol and they are better known in the medical community because of their prevalence. However, RDs such as Spinal Muscular Atrophy, Huntington’s Disease, Neurofibromatosis, and IPEX, which were mentioned by many respondents, have a long diagnostic odyssey, and patients often face long delays in getting the right diagnosis.

Furthermore, the time for diagnosis in India is expected to be highly variable and is highly dependent on social determinants of health such as geographic location (urban vs rural), access to technologies, access to the healthcare, socioeconomic status, scientific knowledge, and awareness of primary care provider. Thus, the responses received are just indicative of the current situation based on respondents’ experience but are not an absolute reflection of the real scenario, as the complete background of the respondents is not well known.

### 3.3 Treatment and diagnosis options

We noted that the perception of HCPs about treatment modalities varies largely with their educational backgrounds. Most respondents selected “symptomatic treatment,” which relates to the real scenario as curative treatment is lacking for most RDs. Globally as well treatment for majority of rare diseases is restricted to palliative care. However, a myriad of different options has been explored. Over the last few decades small molecules, oligonucleotides (including antisense oligos), biologics (including monoclonal antibodies and enzyme replacement therapies, ERTs) and most recently vectorized gene therapies have been approved for over 900 rare disease indications [14]. For rare diseases with monogenic etiologies, gene replacement is an alternative, provided the mutation in the gene does not result in a gain of function. Other mechanisms of action including but not limited to gene editing, exon skipping, and promoter modulation can be used as well to replace function over a threshold. Gene therapy using viral vectors treatment of diseases such as retinapigmentosa, spinal mucosal atrophy are approved globally [15] and others such as hemophilia, sickle cell disease and others are in Phase III trials [16–18]. Table 3 shows a list of rare diseases for which gene therapies are in clinical development. In India most of these treatments are either not available or accessible due to prohibitive costs. Recently a few academic institutes and pharma/biotech companies have initiated development of gene therapies. The GROW lab in Narayana Nethralaya; inSTEM labs; Immuneel Therapeutics, and Intas Biopharma are some of the institutes/hospitals that are working on gene therapies. However extensive work is needed through patient advocacy groups, industry and government to ensure accessibility of these drugs to patients.

**Table 3.**
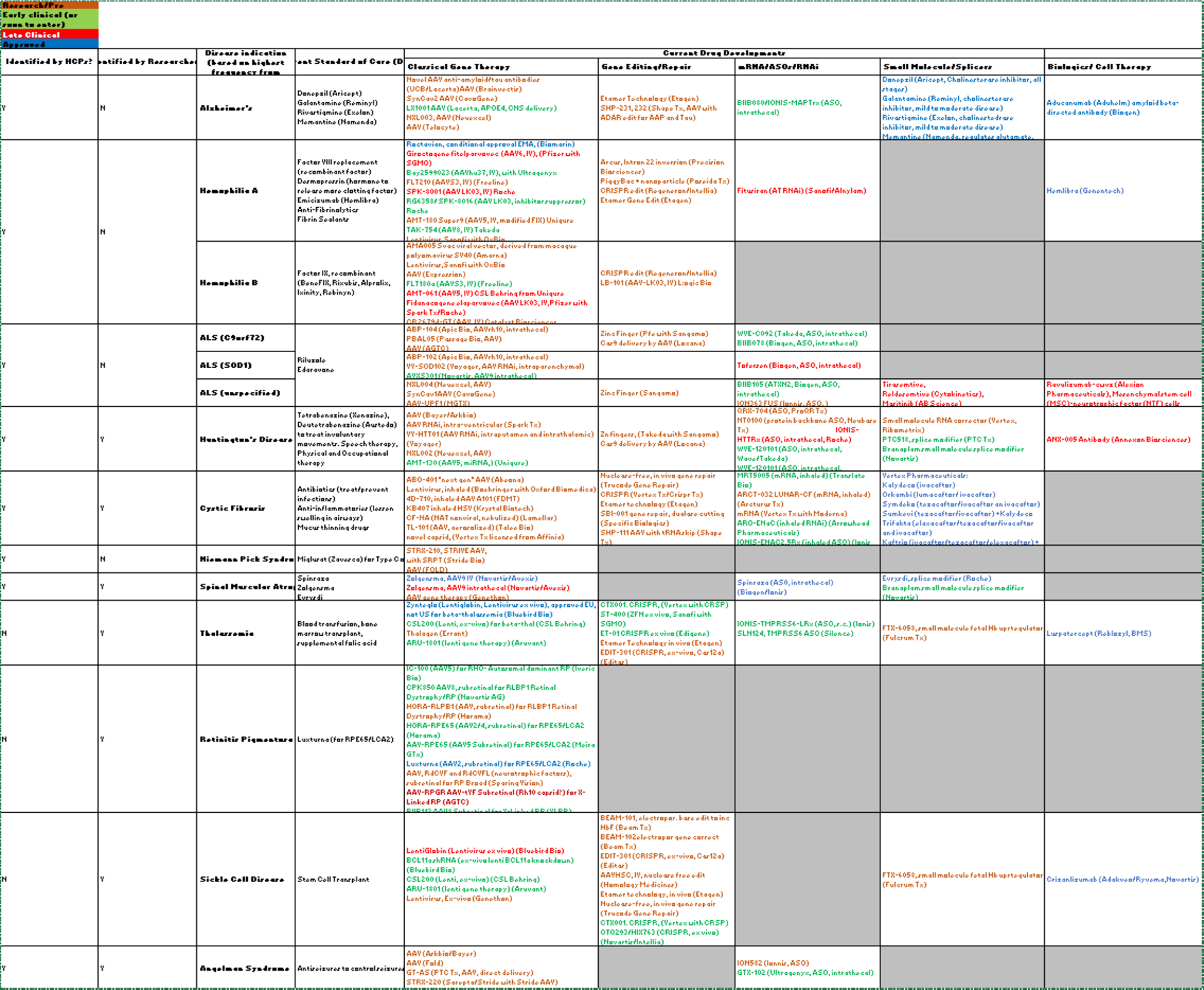
List of rare diseases for which gene therapies are in clinical development

### 3.4 Source of awareness

In the case of RDs, patient groups have been the main drivers for raising awareness about their diseases globally. They have also played a major role in shaping the agenda for RDs and integrating RD education in the medical education system [19]. Most RD patient organizations are actively and successfully involved in numerous activities for and by doctors, researchers, and clinicians specializing in the field of their respective diseases. In India as well, patient organizations have played a significant role in driving the RD policy and creating awareness in general public [9]. The importance of patient advocates in raising awareness needs to be acknowledged by the medical and research fraternity. Further, more formal mechanisms for systematic participation of patient organizations in creating modules for awareness and medical education must be developed. In this survey, 50% of the respondents selected multiple sources for receiving information about RDs, which included research papers, social media, and print media. Among these sources, research papers were cited to be the most preferred source of keeping oneself updated. However, there has been a significant lack of research on RDs in India and focus needs to be given to encourage more research that would improve understanding of disease prevalence and natural history in the Indian population.

### 3.5 Geographical distribution and 3.6 Family History

This question was included in the survey as certain genetic conditions are observed to be more prevalent in certain Indian states. This can be due to social and cultural practices such as consanguinity and endogamy. The southern states have been reported to show a high incidence of metabolic conditions, thalassemia, and other rare genetic diseases which has been correlated to the high prevalence of consanguinity in these populations [20]. Another example is the Agarwal community of Rajasthan, which is seen to have a high prevalence of *MLC1* gene-related Megalencephalic Leukoencephalopathy with subcortical cysts [21, 22]. Hemoglobinopathies such as sickle cell anemia, β-thalassemia, and other thalassemia variants are more prevalent in the Eastern states [23].

A significant percentage of respondents (50% HCPs and 37% researchers) in this survey did not believe that some RDs could be more prevalent in certain regions; however, they believed that family history has a significant role to play in RDs. This shows that although the respondents acknowledge the role of genetics in RDs, they do not necessarily acknowledge the role of genetics at the population level. Also, this indicates that there is a lack of awareness about the studies in India, which have established that some of these diseases indeed have a high prevalence inn certain community.

## 4. Recommendations

Based on our observations from the survey and detailed discussions with various researchers and healthcare workers who work with patients with RDs, we have the following recommendations:

- Multifaceted awareness campaigns that consider existing inequality in the Indian healthcare system must be conducted by government and advocacy groups.
- General awareness among primary HCPs to identify people living with RDs in the population should be increased through various modes, and the identified people should be connected to a proper referral system. Primary HCPs must be empowered with knowledge and infrastructure to screen RDs.
- More targeted training and education, and real-world case studies must be done for HCPs in their specific areas of practice such as neuroscience, immunology, and metabolic diseases.
- Courses on RDs must be integrated in the biological sciences and medical sciences curricula.
- The role of patient organizations and patient advocates in driving awareness about their diseases must be acknowledged. Further, they must be empowered to lead and participate in all kinds of awareness programs including medical education.
- Medical students, baccalaureate and masters students, and researchers should be encouraged to volunteer with patient organizations on internship projects, which will help in sensitizing them with the challenges faced by the RD community [9].

In conclusion, we have surveyed the current level of awareness about RDs in India. We have commented on the perception of HCPs and researchers in India about RD diagnosis and treatment options. Finally, we have made recommendations to further increase the awareness of RDs.

